# Codon-optimized TDP-43-mediated neurodegeneration in a Drosophila model for ALS/FTLD

**DOI:** 10.1101/696963

**Authors:** Tanzeen Yusuff, Shreyasi Chatterjee, Ya-Chu Chang, Tzu-Kang Sang, George R. Jackson

**Affiliations:** Department of Neuroscience and Cell Biology, University of Texas Medical Branch at Galveston, Galveston, TX 77550, USA; Mitchell Center for Neurodegenerative Diseases, University of Texas Medical Branch at Galveston, Galveston, TX 77550, USA; Department of Neurology, University of Texas Medical Branch at Galveston, Galveston, TX 77550, USA; Department of Biochemistry and Molecular Biology, University of Texas Medical Branch at Galveston, Galveston, TX 77550, USA; Department of Biochemistry and Molecular Biology, Pennsylvania State University, University Park, PA 16802, USA; Institute of Biotechnology, Department of Life Science, National Tsing Hua University, Hsinchu, Taiwan; Brain Research Center, National Tsing Hua University, Hsinchu, Taiwan; Department of Neurology, Baylor College of Medicine, Michael E. DeBakey VA Medical Center, Houston, TX 77030, USA; National Parkinson’s Disease Research Education and Clinical Center, Michael E. DeBakey VA Medical Center, Houston, TX 77030, USA

**Keywords:** Drosophila, neurodegeneration, TDP-43, ALS, FTLD

## Abstract

Transactive response DNA binding protein-43 (TDP-43) is known to mediate neurodegeneration associated with amyotrophic lateral sclerosis (ALS) and frontotemporal lobar degeneration (FTLD). The exact mechanism by which TDP-43 exerts toxicity in the brains of affected patients remains unclear. In a novel *Drosophila melanogaster* model, we report gain-of-function phenotypes due to misexpression of insect codon-optimized version of human wild-type TDP-43 (CO-TDP-43) using both the binary GAL4/UAS system and direct promoter fusion constructs. The CO-TDP-43 model showed robust tissue specific phenotypes in the adult eye, wing, and bristles in the notum. Compared to non-codon optimized transgenic flies, the CO-TDP-43 flies produced increased amount of high molecular weight protein, exhibited pathogenic phenotypes, and showed cytoplasmic aggregation with both nuclear and cytoplasmic expression of TDP-43. Further characterization of the adult retina showed a disruption in the morphology and function of the photoreceptor neurons with the presence of acidic vacuoles that are characteristic of autophagy. Based on our observations, we propose that TDP-43 has the propensity to form toxic protein aggregates via a gain-of-function mechanism, and such toxic overload leads to activation of protein degradation pathways such as autophagy. The novel codon optimized TDP-43 model is an excellent resource that could be used in genetic screens to identify and better understand the exact disease mechanism of TDP-43 proteinopathies and find potential therapeutic targets.

## INTRODUCTION

Transactive response DNA binding protein-43 (TDP-43), encoded by *TARDBP* gene in the human genome, has been identified as a major component for the pathology of motor neuron diseases and related neurodegenerative diseases (Neumann *et al.* 2006; Hasegawa *et al.* 2007). TDP-43 is a highly conserved and ubiquitously expressed protein that is primarily involved in regulation of RNA levels, RNA trafficking, and alternative splicing. The presence of tau-negative TDP-43 and ubiquitin-positive inclusion bodies is a major disease hallmark of amylotrophic lateral sclerosis (ALS) and frontotemporal lobar dementia (FTLD) (Neumann *et al.* 2006; Arai *et al.* 2006; Mackenzie and Rademakers 2008; Lee *et al.* 2012). In the diseased state, TDP-43 is found to be ubiquitinated and phosphorylated, and exhibits truncated C-terminal fragments and insoluble inclusions. The distinctive pathology of TDP-43 mediated neurodegeneration also involves its mislocalization to the cytoplasm and the loss of normal nuclear expression (Arai *et al.* 2009; Barmada *et al.* 2010; Guo *et al.* 2011; Lee *et al.* 2012; Nguyen *et al.* 2018). Mutations in the *TARDBP* gene are associated with both familial and sporadic cases of these diseases. Most of the dominant missense mutations are present in the glycine-rich domain near the C-terminal of TDP-43 (Nonaka *et al.* 2009; Lee *et al.* 2012), and have been linked to the formation of toxic TDP-43 aggregates that mediate neurodegeneration (Igaz *et al.* 2011). Protein-protein interactions, hyperphosphorylation, ubiquitination, and cleavage of the prion-like C-terminal fragment have been implicated in the formation of these TDP-43 aggregates (Johnson *et al.* 2009). In addition, the increased load of toxic protein aggregates has been suggested to cause defects in protein degradation systems, including autophagy and the ubiquitin proteasome system (UPS) (Rubinsztein 2006; Blokhuis *et al.* 2013). In order to better understand the pathogenic mechanisms of TDP-43 mediated neurodegeneration, many cellular and animal models have been generated in both vertebrates and invertebrates, which include gain-of-function, RNA interference (RNAi) mediated suppression, and loss-of-function models (Johnson *et al.* 2008; Feiguin *et al.* 2009; Lu *et al.* 2009; Wegorzewska *et al.* 2009; Li *et al.* 2010; Stallings *et al.* 2010; Tsai *et al.* 2010; Estes *et al.* 2011; Gendron and Petrucelli 2011; Vaccaro *et al.* 2012; Romano *et al.* 2012; Choksi *et al.* 2014).

*Drosophila melanogaster* has been widely utilized to study neurodegenerative diseases in an *in vivo* model system (Sang and Jackson 2005). We and others have previously shown that overexpressing toxic proteins such as full-length human tau, alpha-synuclein, or huntingtin in the Drosophila eye or neuromuscular junction results in degenerative phenotypes that are ideal for high-throughput screens, as well as for studying pathogenic mechanisms of the disease (Feany and Bender 2000; Auluck *et al.* 2002; Shulman and Feany 2003; Blard *et al.* 2007; Chatterjee *et al.* 2009; Wegorzewska *et al.* 2009; Li *et al.* 2010; Shulman *et al.* 2014). For example, loss-of-function models generated using deletion, nonsense or null mutations, and RNA-interference mediated knockdown of the Drosophila homolog of TDP-43, TBPH, showed shortened lifespan, locomotor and neuromuscular junction (NMJ) defects, and decreased dendritic branching of DA neurons (Feiguin *et al.* 2009; Lu *et al.* 2009). Furthermore, gain-of-function transgenic fly models overexpressing disease-specific variants of human TDP-43 (hTDP-43) showed decreased longevity, decreased locomotor activity, and increased morphological defects of motor neurons, along with axonal damage and, in some cases, neuronal loss (Lu *et al.* 2009; Li *et al.* 2010, 2011; Hanson *et al.* 2010; Ritson *et al.* 2010; Voigt *et al.* 2010; Estes *et al.* 2011; Guo *et al.* 2011; Miguel *et al.* 2011; Langellotti *et al.* 2016; Chang and Morton 2017; Pons *et al.* 2017). These gain-of-function mutations only account for about 10% of familial cases of ALS/FTLD, while 90% of affected individuals are sporadic cases involving wild-type TDP-43 mediated neurodegeneration (Nguyen *et al.* 2018). However, current studies involving wild-type TDP-43 have reported only subtle phenotypes that were difficult to quantify or did not exhibit robust disease-associated pathology. Therefore, there is a need for a robust model of wild-type TDP-43 mediated pathology to understand the cellular mechanisms associated with ALS/FTLD.

We generated an overexpression model of the human wild-type TDP-43 transgene by codon-optimization to accommodate insect transcriptional and translational machinery. It has been shown, even in *Drosophila melanogaster*, that certain 3-base pair sequences or codons in the mRNA transcript are more optimal in translating into the same amino acid over others (Powell and Moriyama 1997; Welch *et al.* 2009). Using this phenomenon, we manipulated the human *TARDBP* gene by altering the coding region so that codons were optimized in a *Drosophila melanogaster* cellular environment to maximize TDP-43 expression, henceforth referred to as CO-TDP-43. In contrast to previous fly models, we demonstrate that the CO-TDP-43 lines lead to increased TDP-43 expression and form toxic cytoplasmic aggregates that gives rise to strong phenotypes when expressed in the fly retina, wing, and notum. Further characterization of the retinal phenotype revealed a disruption in the internal morphology and function of the photoreceptor neurons, as well as presence of acidic autophagic-lysosomal vacuoles that are positive for key autophagy proteins. Our CO-TDP-43 model recapitulates phenotypes of ALS/FTLD disease pathology and is an ideal resource for investigating the mechanisms of pathogenesis for these diseases.

## MATERIALS AND METHODS

### Fly stocks and genetics

Codon optimized TDP-43 gene was synthesized from DNA2.0 (ATUM, Newark, CA, USA). The complete sequence of the codon optimized TDP-43 is provided in the **Supplemental Fig. S2**. Drosophila kozak sequence (ATCAAC) was added upstream of the start codon for the TDP43 gene. These constructs were subcloned into the Not1-Xba1 site of the modified fly upstream activation sequence (UAS) expression (pEx-UAS) and glass (pEx-gl) vectors (Exelixis, San Francisco, CA, USA). The expression vectors containing the codon optimized TDP-43 gene were then microinjected into the flies to obtain transgenic flies (BestGene, Chino Hills, CA, USA). The expression of non-CO-TDP-43 is driven in the fly eye by the *glass* multimer reporter, GMR-GAL4 on the X-chromosome (Freeman 1996). All transgenic lines, both codon optimized and non-codon optimized, express human wild-type TDP-43. Flies expressing human codon wild-type TDP-43 using the UAS promoter were obtained from Dr. Fen-Biao Gao (University of Massachusetts, Worcester, MA, USA) (Lu *et al.* 2009). SevEP-GAL4 driver (expressed in R7 and R8 photoreceptor neurons) was recombined with UAS-TDP-43CO to obtain stable transgenic flies expressing w1118;SevEP-GAL4,UAS-TDP-43CO/CyO;+. The GMR-GAL4 on the X-chromosome was placed in trans to the gl-TDP-43CO line to generate GMR-GAL4;gl-TDP-43CO/CyO transgenic flies. The following stocks were obtained from Bloomington Drosophila Stock Center (Bloomington, Indiana University, IN, USA): w1118;UAS-LacZ, w1118,GMR-myr-mRFP, y1,w1118;Sp/CyO;eGFP-ATG5, y1,w1118;UASp-GFP-mCherry-ATG8, GMR-GAL4(X) (eye specific), w1118;*SevEP*-GAL4 (R7 and R8 in photoreceptor cells), y1,w1118;*Rh1*-GAL4/CyO (expressed in R1-R6 photoreceptor cells), w1118,*beadex*^MS1096^-GAL4 (wing driver), w1118;*Scabrous*-GAL4 (sensory organ precursor and wing discs driver), and y1,w*; CCAP-GAL4 (driver expressed in CCAP/bursicon neurons in ventral nerve cord and subesophageal ganglion in adult brain). *Eq*-GAL4 (bristle driver) was obtained from Dr. Hugo J. Bellen (Baylor College of Medicine, Houston, TX). All crosses were set and flies were maintained at room temperature (22°C) in standard D. melanogaster Jazzmix medium (Applied Scientific, Fisher Scientific, Pittsburgh, PA, USA).

### Immunohistochemistry

Adult retina and imaginal eye discs from third instar larvae were dissected and fixed in 4% paraformaldehyde for 1 hour on ice. Adult retina was washed in 0.5% PTX for 3 hours to reduce autofluorescence. The tissues were blocked in 0.8% PBS+Triton-X+BSA for 2 hours and incubated with primary antibody overnight at 4°C. The tissues were incubated in secondary antibody for 2 hours at room temperature, washed in 0.1% PBS+Triton-X and mounted on glass slides with Vectashield (Vector Laboratories, Burlingame, CA, USA). Tissues were stained with the following antibodies: mouse monoclonal anti-TDP-43 antibody (1:500, Abcam, Cambridge, MA, USA), rabbit polyclonal anti-TDP-43 antibody (1:500, Proteintech, Chicago, IL,USA), rat monoclonal anti-Elav (1:20, DSHB, University of Iowa, Iowa City, IA, USA), mouse monoclonal anti-GFP (1:400, Millipore, Billerica, MA, USA), Alexa Fluor 633-conjugated Phalloidin (1:30, Invitrogen, Grand Island, NY, USA), Alexa Fluor 488 conjugated chicken anti-rat (1:400, Invitrogen, Grand Island, NY, USA) Alexa Fluor 568 conjugated goat anti-rabbit (1:400, Invitrogen, Grand Island, NY, USA) and Alexa Fluor 568 conjugated goat anti-mouse (1:400, Invitrogen, Grand Island, NY, USA).

### Immunoblotting

Overexpression of CO-TDP-43 in the fly eye was used to measure total protein levels by immunoblotting. Approximately 50 fly heads were decapitated and homogenized for 1 min in homogenization buffer (10mM Tris-HCl, 0.8 M NaCl, 1 mM EGTA, pH 8.0 and 10% sucrose) along with 1X PhosSTOP phosphatase and 1X cOmplete protease buffer (Roche Applied Science, Indianapolis, IN, USA). The homogenized samples were centrifuged at 4°C for 15 min at 18,000g. The supernatant was collected and equal parts of the supernatant and Laemmle sample loading buffer with β-mercaptoethanol (Bio-Rad, Hercules, CA, USA) was added for each sample. Following a brief pulse centrifugation, samples were loaded on 4-20% SDS-PAGE gels (Bio-Rad, Hercules, CA, USA) for electrophoresis. For higher molecular weight species detection, the fly heads were homogenized in 1X PBS along with the same protease and phosphatase inhibitors. Non-reducing sample loading buffer (Nupage sample buffer, Life Sciences, Grand Island, NY, USA) was added to the supernatant without β-mercaptoethanol. The blots were blocked in 5% milk, incubated with primary antibodies overnight at 4°C, washed in 1X TBS+Tween, and incubated with secondary antibody for 1 hour at room temperature. The following antibodies were used: mouse monoclonal anti-TDP-43 antibody (1:1000, Abcam, Cambridge, MA, USA), mouse monoclonal anti-tubulin antibody (1:1000, DSHB, University of Iowa, Iowa City, IA, USA) and secondary anti-mouse IgG-HRP (1:2000, GE Healthcare).

### Lysotracker Staining

For LysoTracker staining, imaginal eye discs from the third instar larvae were dissected in 1X PBS solution without fixative. The eye discs were then stained with 100 nM LysoTracker Red DND-99 (Invitrogen) for 2 minutes, followed by a 1 minute wash in 1X PBS. The tissues wer mounted on a glass slide with a drop of 1X PBS solution; no Vectashield was added. The coverslip was sealed with nail polish and visualized immediately using a confocal microscope. The z-stack images were analyzed using the ImageJ software (Schneider *et al.* 2012).

### Electroretinogram

ERG was recorded in 1 day old flies using the same methods as previously described (Fabian-Fine *et al.* 2003; Williamson *et al.* 2010). Briefly, flies were glued on glass slides using Elmer’s non-toxic glue. Both the reference and recording electrodes were made of glass pipettes filled with 3M KCl. The light stimulus was computer-controlled using white light-emitting diode system (MC1500; Schott), and was provided in 1-s pulses. The data was recorded using Clampex software (version 10.1; Axon Instruments) and measured and analyzed using Clampfit software (version 10.2; Axon Instruments).

### Microscopy

The adult eye, wing and bristle pictures were taken with a Nikon AZ100M microscope equipped with a Nikon DS-Fi1 digital camera (Nikon Instruments, Melville, NY, USA). Extended depth of focus (EDF) and volumetric images were taken using the Nikon NIS-Elements AR 3.0 software as previously described (Ambegaokar and Jackson 2011). The scanning electron microscope (SEM) images were taken using JSM-6510LV SEM (JEOL USA, Peabody, MA, USA). The confocal images were taken with a Zeiss LSM 510 UV META laser scanning confocal microscope using 40X water and 63X oil-immersion high-resolution objectives. These images were analyzed using the LSM Image Browser and NIH ImageJ software (Schneider *et al.* 2012).

### Statistical Analysis

Quantification of LysoTracker staining was performed using the NIH Image J software (Schneider *et al.* 2012). The measurements and histograms represent mean±SEM and plotted using Microsoft Excel and SigmaPlot (version 10.1) software. Statistical analysis was performed using one-way ANOVA with Bonferroni’s correction and paired Student’s t-test with two-tailed distributions of equal variance.

### Data Availability

The codon-optimized TDP-43 fly lines are available upon request.

## RESULTS

### Codon-optimized wild-type TDP-43 flies exhibit an age-dependent robust eye phenotype

We generated multiple codon-optimized CO-TDP-43 transgenic fly lines to investigate TDP-43 mediated neurodegeneration. We utilized both the yeast GAL4/UAS binary system (Brand and Perrimon 1993) and a *glass (gl)* promoter direct fusion construct specifically generated to study TDP-43 mediated effects on the fly retina (Fig. 1A-B). In addition, we also used another eye promoter, *Sevenless* (*SevEP*-GAL4), that only expresses in a subset of photoreceptor neurons (R7 and R8) and cone cells (Therrien *et al.* 1999). To highlight the robust effect observed in our CO-TDP-43 lines, we compared the phenotypes to a previously reported human TDP-43 transgenic line, which we denote as non-CO-TDP-43 (Lu *et al.* 2009; Choksi *et al.* 2014).

**Figure 1.**
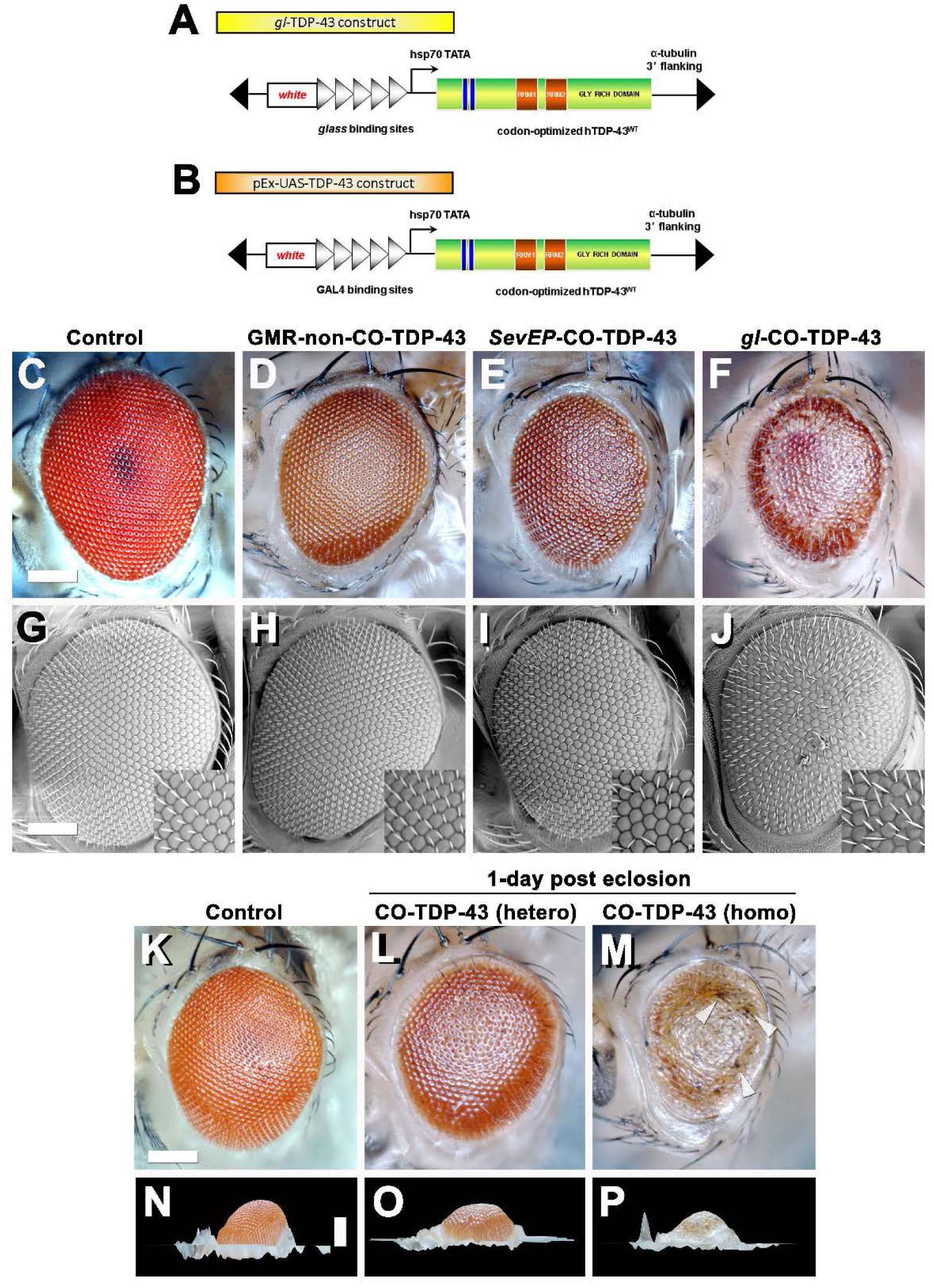
Misexpression of codon optimized TDP-43 induces depigmentation and irregularities in bristles compared to existing human wild-type TDP-43 lines. Transgenic flies stably express wild-type or codon optimized TDP-43 using eye promoters. All stocks were kept and maintained at room temperature (22°C). **(A-B)** Schematic of the codon optimized TDP-43 constructs using *glass* direct fusion promoter vector and UAS vector used to create the transgenic codon optimized TDP-43 lines. **(C-J)** Photomicrograph and scanning electron microscopy (SEM) images of the adult retina at 10 days post-eclosion (scale bar 50 uM). Compared to existing human wild-type TDP-43 transgenic flies expressed using GMR-GAL4 promoter **(D and H)**, the codon optimized wild-type TDP-43 flies using *gl*ass direct fusion promoter **(F and J)** exhibit a robust eye phenotype including depigmentation, disruption in planar polarity and loss of bristles. *SevEP*-GAL4, a selective R7 and R8 photoreceptor neuron driver, recombined with codon optimized wild-type TDP-43 **(E and I)** also shows the same phenotypes. **(C and G)** are controls. **(K-P)** The robust phenotype mediated by codon optimized TDP-43 is both age and dosage dependent. At 1 day post-eclosion, codon optimized TDP-43 **(L)** shows less depigmentation compared to 10 days post-eclosion **(E)**. A homozygous codon optimized TDP-43 expression **(M)** shows a dramatically more robust phenotype with some necrosis (white arrowheads) at 1 day post-eclosion. Compared to control flies **(K and N)**, both hetero- and homozygous codon optimized TDP-43 shows decreased volume **(O and P respectively)**. Scale bar: 100 nm. Genotypes: **(C and G)** Canton S, **(D and H)** w^1118^/+;GMR-GAL4/+;UAS-hTDP-43^WT^/+, **(E and I)** w^1118^/+;*SevEP*-GAL4,UAS-TDP-43^CO^/+;+, (F and J) w^1118^/+;*gl*-TDP-43^CO^/+;+, **(K and N)** Canton S, **(L and O)** w^1118^/+;*gl*-TDP-43^CO^/+;+, **(M and P)** w^1118^;*gl*-TDP-43^CO^;+.

Heterozygous expression of CO-TDP-43 using the *gl* promoter caused depigmentation, roughness, disruption of polarity, and loss of inter-ommatidial bristles (Fig. 1F and J). The CO-TDP-43 expressed using *SevEP*-GAL4 showed a similar but milder phenotype of the eye (Fig. 1E and I). In comparison to the CO-TDP-43 flies, the non-CO-TDP-43 transgenic flies (Fig. 1 D and H) did not show a robust eye phenotype and appeared to be similar in morphology to the wild-type control flies (Fig. 1C and G). Interestingly, the eye phenotype observed with heterozygous *gl*-CO-TDP-43 flies were age dependent. At day-1 post-eclosion, CO-TDP-43 exhibited a mild phenotype (Fig. 1L) that worsened by day 10 (Fig. 1M). In contrast, flies with two copies of the CO-TDP-43 transgene showed a strong phenotype at day-1 post-eclosion, with apparent necrotic patches or hyperpigmentation that worsened with age (Fig. 1M, white arrowheads). In addition, the CO-TDP-43 flies with either one or two copies of the TDP-43 transgene showed less eye volume than wild-type control flies at day-1 post-eclosion (Fig. 1O and P). We also found that overexpression of CO-TDP-43 using GMR-GAL4 driver led to pupal lethality at 18°C and 25°C (**Supplemental Table S1**), with some escapers at 18°C that showed necrotic patches (**Supplemental Fig. S1**). Taken together, our results showed that CO-TDP-43 transgenic flies have a more robust eye phenotype indicative of neurodegeneration in retinal cells compared to non-CO-TDP-43 transgenic flies.

### Misexpression of codon-optimized wild-type TDP-43 leads to necrosis and severe phenotypes in wings and notum

TDP-43 associated pathology in ALS patients have been linked to significant neuronal loss and early axonal atrophy in sensory nerves (Heads *et al.* 1991; Mochizuki *et al.* 2011). For example, Vaughan and colleagues reported that the pathogenic A315T mutation in TDP-43 affects neurite growth and decreased dendritic branching of sensory neurons (Vaughan *et al.* 2018). In fact, previously reported Drosophila neurodegeneration models showed that overexpression of the neurotoxic ataxin-1 mutant in sensory precursors using the *scabrous*-GAL4 (*sca*-GAL4) driver leads to loss of bristles in the adult fly (Tsuda *et al.* 2005). Similarly, we previously showed that misexpression of fly dVAP33, a gene linked to ALS, using *sca*-GAL4 leads to loss of notal macrochaetae (Ratnaparkhi *et al.* 2008). To further investigate the phenotypic effects of CO-TDP-43 on sensory precursor cells of the wing and notum, we used multiple wing and bristle drivers to misexpress TDP-43 protein, including *beadex^MS1096^*-GAL4 (*bx^MS1096^*-GAL4), *sca*-GAL4, *equate*-GAL4 (*eq*-GAL4), and CCAP-GAL4. We found that non-CO-TDP-43 transgene expressed using *bx^MS1096^*-GAL4 led to viable adults with shriveled wings, with some flies having wings that were either necrotic or had areas of hyperpigmentation (Fig. 2B). In contrast, CO-TDP-43 flies using the same driver exhibited a more severe phenotype, with pharate adults and very small and severely malformed wings with necrotic or hypermelanized patches (Fig. 2C). Interestingly, unlike the previously reported model of ALS, neither non-CO-TDP-43 nor CO-TDP-43 had any effect on macrochaetae (bristles) on the notum when misexpressed using *sca*-GAL4 driver. Instead, the CO-TDP-43 transgenic flies produced pharate adults with necrotic wings that were unable to expand (Fig. 2F).

**Figure 2.**
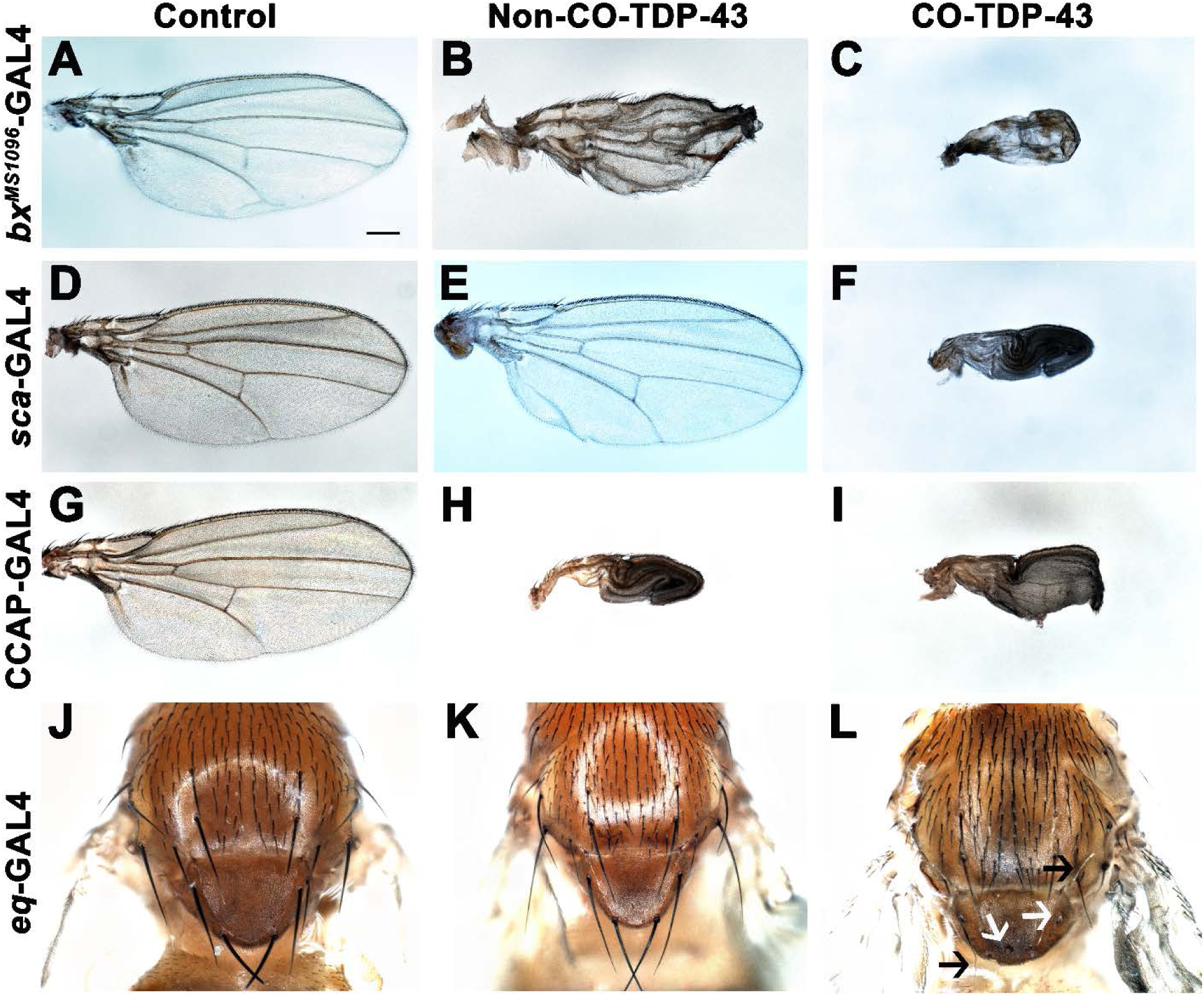
Misexpression of codon optimized TDP-43 leads to wing expansion and swelling defects as well as singed and loss of bristles in the fly notum. **(A-C)** Codon optimized TDP-43 expressed in the wings using the wing-specific driver *bx^MS1096^*-GAL4 leads to pharate adults with smaller, swollen and necrotic wings **(C)**, as compared to healthy, viable adults expressing human wild-type TDP-43 with crumpled wings **(B)** and normal wings with the driver alone **(A)**. **(D-F)** Using *sca*-GAL4 driver, human wild-type TDP-43 flies **(E)** have normal wings, similar to controls with the driver alone **(D)**, while codon optimized TDP-43 causes pharate adults with smaller, necrotic wings with expansion defect **(F)**. **(G-I)** CCAP-GAL4, expressed in CCAP/bursicon neurons in the ventral nerve cord and the subesophageal ganglion in the adult brain, driven expression of codon optimized TDP-43 **(I)** as well as human wild-type TDP-43 **(H)** also exhibit similar wing expansion defects, compared to the driver alone **(G)**. **(J-L)** A bristle specific driver, *eq*-GAL4, causes a dramatic loss of bristles (white arrows) and singed bristles (black arrows) with codon optimized TDP-43 flies **(L)**, while human wild-type TDP-43 **(K)** and driver alone **(J)** develop normal bristles. Scale bar: 200um. Genotypes: **(A)** w^1118^,*bx^MS1096^*-GAL4/+;+;+, **(B)** w^1118^,*bx^MS1096^*-GAL4/+;+;UAS-hTDP-43^WT^/+, **(C)** w^1118^,*bx^MS1096^*-GAL4/+;+;UAS-TDP-43^CO^/+, **(D)** w^1118^/+;*sca*-GAL4/+;+, **(E)** w^1118^/+;*sca*-GAL4/+;UAS-hTDP-43^WT^/+, **(F)** w^1118^/+;*sca*-GAL4/+;UAS-TDP-43^CO^/+, **(G)** y^1^,w*/+;CCAP-GAL4/+;+, (H) y^1^,w*/+;CCAP-GAL4/+;UAS-hTDP-43^WT^/+, **(I)** y^1^,w*/+;CCAP-GAL4/+;UAS-TDP-43^CO^/+, **(J)** w^1118^/+; *eq*-GAL4/+;+;+, **(K)** w^1118^/+; *eq*-GAL4/+;+;UAS-hTDP-43^WT^/+, **(L)** w^1118^/+; *eq*-GAL4/+;+; UAS-TDP-43^CO^/+.

Since we failed to see an effect of TDP-43 on macrochaetae using *sca*-GAL4, we used another bristle-specific driver, *eq*-GAL4, to misexpress non-CO and CO-TDP-43 in the fly notum (Tang and Sun 2002). While both control and non-CO-TDP-43 flies showed normal macrochaetae formation (Fig. 2J and K), CO-TDP-43 showed a dramatic loss or defective notal macrochaetae (Fig. 2L). Furthermore, Vanden Broeck and colleagues previously showed that both up and downregulation of fly dTDP-43 cause selective apoptosis in the crustacean cardioactive peptide (CCAP)/bursicon neurons (Vanden Broeck *et al.* 2013). Loss of CCAP/bursicon neurons have been shown to cause pupal lethality with escapers that show wing expansion defect phenotypes (Park *et al.* 2003). Upon expression of non-CO-TDP-43 in the CCAP/bursicon neurons using CCAP-GAL4, we observed a similar wing expansion defect in adults (Fig. 2H). Misexpression of CO-TDP-43 in CCAP/bursicon neurons resulted in smaller, necrotic and swollen wings compared to control flies (Fig. 2I). In summary, these results suggest that misexpression of CO-TDP-43 in flies leads to smaller wings with abnormal morphology and macrochaetae irregularities compared to misexpression of non-CO-TDP-43.

### Increased expression of codon-optimized TDP-43 exhibits disease-specific cytoplasmic mislocalization and aggregation

The robustness of the external phenotypes observed with CO-TDP-43 prompted us to examine the protein expression levels of the TDP-43 transgene in these flies. We next used multiple *gl* direct fusion CO-TDP-43 lines as well as the recombined *SevEP*-GAL4 line to examine TDP-43 expression levels in the fly eye. Compared to the GMR-GAL4 driven non-CO-TDP-43 transgenic flies, the CO-TDP-43 flies showed a 2-fold increase in monomeric total TDP-43 protein in multiple lines (Fig. 3A, lane 5, 6, 7 and 8). The line showing the highest increase in protein levels, one of the *gl* direct fusion CO-TDP-43 lines (Fig. 3A, lane 5), also demonstrated a robust eye phenotype (Fig. 1) and was therefore used in subsequent experiments. Predictably, compared to the *gl*-CO-TDP43 line, the recombinant line using *SevEP*-GAL4 to overexpress CO-TDP-43 did not show an increase in total TDP-43 expression, since it is only expressed in a subset of photoreceptor neurons (Fig. 3A, lane 8).

**Figure 3.**
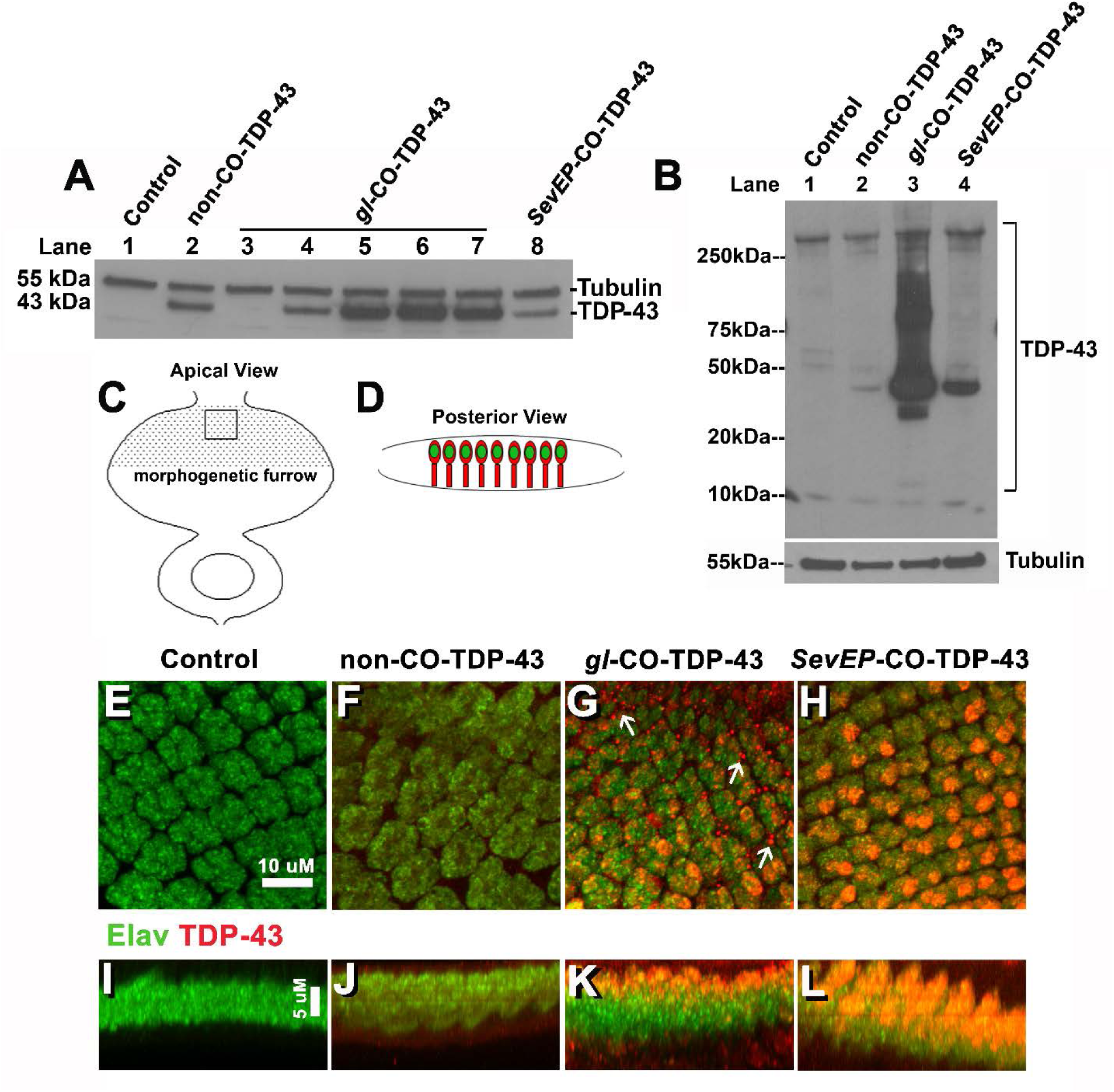
Codon optimized TDP-43 transgenic fly expresses higher protein levels, causing robust mislocalization to the cytoplasm and aggregate formation in larval eye discs. **(A)** Western blot analysis comparing the total TDP-43 levels in eyes from human wild-type TDP-43 flies (lane 2) to the different generated codon optimized TDP-43 lines (lane 3-8) and control flies (lane 1). Among the different codon optimized TDP-43 lines, *gl*-TDP-43^CO3^, *gl*-TDP-43^CO4^ and *gl*-TDP-43^CO5^ (lane 5, 6 and 7 respectively) have the highest expression of TDP-43, almost a 2-fold increase compared to human wild-type TDP-43. Codon optimized TDP-43 expression driven with a selective R7 and R8 photoreceptor neuron driver, SevEP-GAL4, did not show increased expression of the total protein (lane 8). β-tubulin is presented as a loading control. **(B)** Codon optimized TDP-43 (*gl*-TDP-43^CO3^, lane 3) exhibits higher molecular weight species of TDP-43 as well as the known 15kD truncated c-terminal fragment compared to human wild-type TDP-43 (lane 2) or the SevEP-GAL4 driven codon optimized TDP-43 (lane 4). Lane 1 is wild-type control. β-tubulin is presented as a loading control. **(C and D)** Schematic of the third instar larval imaginal eye-antennal disc from an apical and posterior view, respectively. The area represented with the rectangular box in **(C)** is the area imaged. **(E-L)** Confocal images of the third instar imaginal eye discs stained with neuronal marker Elav (green) and TDP-43 (red). There is a greater expression of both nuclear and cytoplasmic TDP-43 in codon optimized lines **(G and H)** as compared to human wild-type TDP-43 **(F)**. The *gl*-TDP-43^CO3^ flies exhibit a more robust mislocalization and aggregation of cytoplasmic TDP-43; white arrows in **(G)**. **(E)** shows the control (scale bar 10 µm). **(I-L)** represents the posterior view of the eye discs to show nuclear and cytoplasmic TDP-43 expression (scale bar 5 µm). Genotypes: **(A)** w^1118^;+;+, w^1118^;GMR-GAL4/+;UAS-hTDP-43^WT^/+, w^1118^;*gl*-TDP-43^CO1^/+;+, w^1118^;+;*gl*-TDP-43^CO2^/+, w^1118^;*gl*-TDP-43^CO3^/+;+, w^1118^;*gl*-TDP-43^CO4^/+;+, w^1118^;*gl*-TDP-43^CO5^/+;+, w^1118^;SevEP-GAL4,UAS-TDP-43^CO^/+;+ (lane 1-8, respectively). **(B)** w^1118^;+;+, w^1118^;GMR-GAL4/+;UAS-hTDP-43^WT^/+, w^1118^;*gl*-TDP-43^CO^/+;+, w^1118^;SevEP-GAL4,UAS-TDP-43^CO^/+;+ (lane 1-4, respectively). **(E and I)** Canton S, **(F and J)** w^1118^;GMR-GAL4/+;UAS-hTDP-43^WT^/+, **(G and K)** w^1118^;*gl*-TDP-43^CO^/+;+, **(H and L)** w^1118^;SevEP-GAL4,UAS-TDP-43^CO^/+;+.

Similar to patients with ALS and FTLD, high-molecular weight toxic species of TDP-43 have been detected in transgenic flies overexpressing TDP-43 containing pathogenic variants (Miguel *et al.* 2011; Chang and Morton 2017). For example, we previously reported high-molecular weight species of TDP-43 in flies overexpressing disease-associated TDP-43 Q331K mutations (Choksi *et al.* 2014). To investigate if we are able to detect these high-molecular weight oligomeric species in our codon optimized lines, we used the *gl* direct fusion line and the stable recombinant *SevEP*-GAL4 driven CO-TDP-43 lines. As expected, we detected higher molecular weight species in SDS-PAGE under non-denaturing conditions in both lines tested, with increased levels in the *gl* driven CO-TDP-43 line that were absent in the non-CO-TDP-43 flies. We also observed 15 kD and 35 kD truncated fragments in the codon optimized flies (Fig. 3B). These bands represent the previously reported caspase cleaved C-terminal fragment that is considered to be the toxic component of TDP-43 aggregates (Liu *et al.* 2014; Chiang *et al.* 2016).

The distinctive pathology of TDP-43 mediated neurodegeneration involves its mislocalization to the cytoplasm and loss of normal nuclear expression (Neumann *et al.* 2006; Lee *et al.* 2012). Therefore, we further investigated the localization of CO-TDP-43 in neuronal cells. When co-stained with *Elav* and TDP-43, the eye discs showed higher nuclear and cytoplasmic expression of TDP-43 in both CO-TDP-43 lines (*gl* and *SevEP*-GAL4 driven) compared to non-CO-TDP-43 flies. In particular, the *gl* driven CO-TDP-43 flies showed a more robust mislocalization of TDP-43 in the cytoplasm along with cytoplasmic aggregates (Fig. 3G, 3I-L). Overall, these observations indicate that a higher level of TDP-43 protein has the propensity to form protein aggregates via a gain-of-function mechanism, similar to other neurodegenerative proteins such as tau, aβ, and alpha-synuclein.

### Morphological and functional disruption of photoreceptor neurons induced by codon-optimized TDP-43

Based on severe retinal phenotypes observed with toxic aggregates of TDP-43 protein, we further investigated the internal cellular morphology of the photoreceptor neurons. We utilized another eye-specific driver, *Rh1*-GAL4, which is expressed in R1–R6 neurons starting in late pupal stage and persisting throughout adulthood (Chyb *et al.* 1999). Unlike the GMR-GAL4, *SevEP*-GAL4 or *gl* direct fusion lines, this driver allowed us to examine adult onset expression of TDP-43. In day-7 post-eclosion CO-TDP-43 flies, we observed a degenerative phenotype in the adult retina marked by the loss of rhabdomere structures and vacuolization compared to the control flies (Fig. 4B). Comparatively, using the *gl* direct fusion CO-TDP43 line, we observed the degenerative phenotype as early as day-1 in post-eclosion flies. The *gl*-CO-TDP-43 flies exhibited an altered morphology of the photoreceptor neurons, which appeared to be flattened and had a disruption in rhabdomere separation (Fig. 4D) when visualized in the tangential view of the adult retina. Examination of the longitudinal view of the adult retina showed a marked reduction in thickness and shorter photoreceptor length compared to controls (marked by white lines in Fig. 4E and I). In addition, we found that these photoreceptor neurons were accompanied by large vacuolar structures, and co-staining with *Elav* revealed that TDP-43 was localized both in the nucleus and in the cytoplasm (Fig. 4I-L).

**Figure 4.**
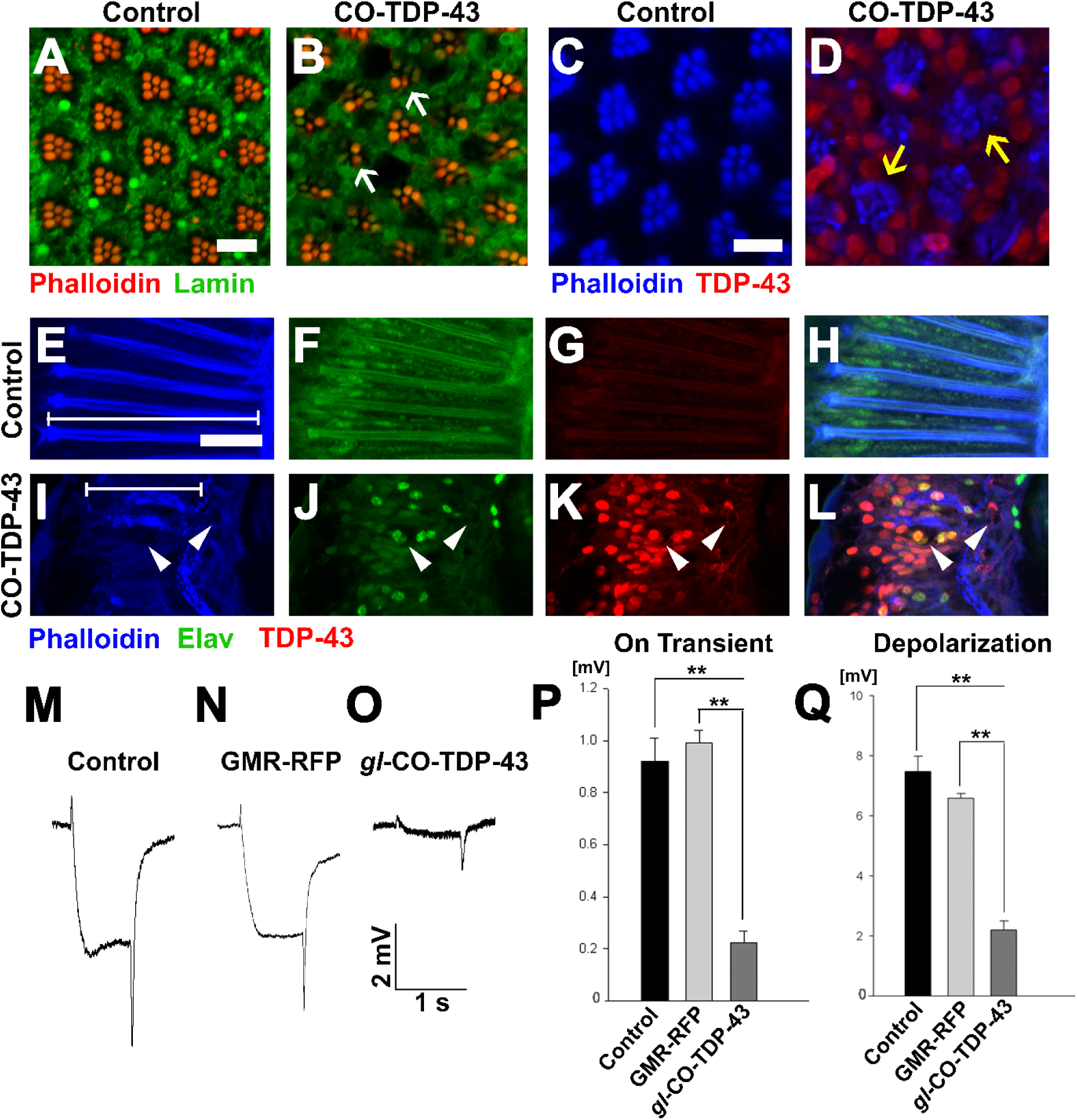
TDP-43 misexpression in adult retina causes degeneration and altered morphology of the photoreceptor neurons. **(A)** UAS-LacZ and **(B)** UAS-TDP-43^CO^ expressed in the retina using *Rh1*-GAL4 that selectively expresses TDP-43 in R1-R6 photoreceptor neurons during the late pupal stage. Compared to control, the codon optimized TDP-43 shows loss of rhabdomeres (white arrows) and degeneration in the 7 days post-eclosion adult retina (scale bar 5 µm). The *gl*-TDP-43^CO3^ flies **(D)** exhibit rhabdomere separation defect and flattened structures of the rhabdomeres (yellow arrows) in 1 day post-eclosion adult compared to GMR-GAL4 control **(C)**, as seen in the tangential view of the retina (scale bar 5 µm). Similarly, in the longitudinal view, the *gl*-TDP-43^CO3^ flies **(I-L)** show altered photoreceptor morphology that appear to be shorter (white lines in **E** and **I**) compared to control **(E-H)**. The codon optimized TDP-43 flies also contain large vacuoles (white arrowheads) in 1 day post-eclosion adult retina (scale bar 10 µm). **(M-O)** ERG traces of wild-type control, GMR-RFP control and *gl*-TDP-43^CO3^ 1 day post-eclosion adults, respectively, are shown. Quantification of the ERG response amplitude for on transient **(P)** and depolarization **(Q)**, along with the traces, show that codon optimized TDP-43 flies have decreased responses for both measures. For on transient effect, n=15 and p< 0.001 between both groups **(P)**, and for depolarization effect, n=15 and p< 0.001 between both groups **(Q)**. Genotypes: **(A)** w^1118^/+;*Rh1*-GAL4/+;UAS-LacZ/+, **(B)** w^1118^/+;*Rh1*-GAL4/UAS-TDP-43^CO^;+, **(C)** Canton S, **(D)** w^1118^;*gl*-TDP-43^CO^/+;+, **(E-H)** Canton S, **(I-L)** w^1118^;*gl*-TDP-43^CO^/+;+, **(M)** Canton S, **(N)** w^1118^;GMR-GAL4/+;UAS-RFP/+, **(O)** w^1118^;*gl*-TDP-43^CO^/+;+.

To investigate the physiological functions of these photoreceptor neurons, we used electroretinogram (ERG) recordings to measure the functionality of active photoreceptor neurons by measuring their response to light stimulus (Dolph *et al.* 2011). In fact, Drosophila neurodegeneration models overexpressing tau and alpha-synuclein exhibited degenerative pathology in the fly retina along with neuronal dysfunction, as detected by ERG recording (Chouhan *et al.* 2016). Using this technique, we investigated whether CO-TDP-43 misexpression affects neuronal functionality compared to wild-type and GMR-RFP controls (Fig. 4M-O). The *gl*-TDP-43^CO3^ flies demonstrated a reduction in both the amplitude of ERG in “on transient” and evoked depolarization at day-1 post-eclosion (Fig. 4P and Q). These effects were not observed with either control. Taken together, these results strongly suggest that CO-TDP-43 misexpression causes structural and functional degenerative phenotypes in the adult retina.

### Codon-optimized wild-type TDP-43 misexpression disrupts cellular lysosomal and autophagic processes

An upregulation of autophagy has been implicated in many neurodegenerative diseases, including ALS (Wong and Cuervo 2010; Brady *et al.* 2011; Sasaki 2011). Therefore, the presence of large vacuolar structures in the adult retina of CO-TDP-43 flies (see Fig. 4) led us to investigate if these vacuoles could possibly be a representation of autophagic intermediates. We performed live imaging of larval eye discs using LysoTracker to detect lysosomes and other acidic organelles, such as autophagosomes, that typically increase in number and/or size during the later stages of autophagy. Upon CO-TDP-43 misexpression, we detected significantly larger number of acidic punctae compared to control flies (Fig. 5C, white arrows and Fig. 5D), suggesting increases in autophagosomes due to elevated levels of TDP-43.

**Figure 5.**
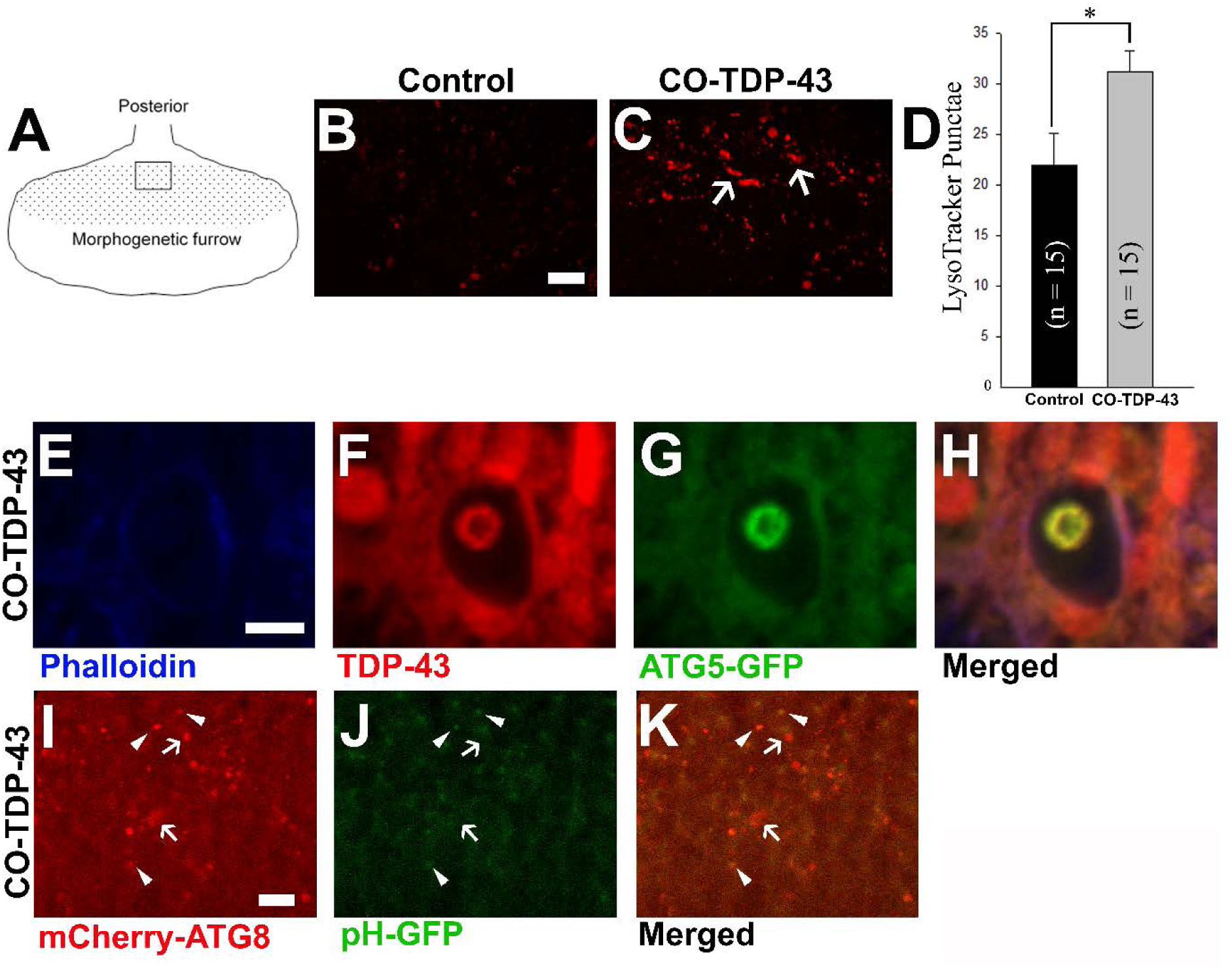
TDP-43 misexpression leads to increased lysosomal vacuoles positive for autophagy markers. **(A)** Schematic of the third instar larvae and the area imaged. Live staining of the LysoTracker dye shows increases in lysosomal puntae in *gl*-TDP-43^CO3^ flies **(C)** compared to control **(B)**. Scale bar equals 10 µm. **(D)** shows quantification of **(B and C)**, n=15 and p=0.02. **(E-H)** Coexpression of *gl*-TDP-43^CO3^ and the autophagic protein Atg5-GFP shows that the large vacuoles present in the 1 day post-eclosion adult retina are positive for Atg5 (scale bar 5 µm). **(I-K)** Another autophagic marker was coexpressed with *gl*-TDP-43^CO3^, Atg8-mCherry-GFP, which is pH sensitive and only expresses GFP at a higher pH content. The TDP-43 expressed flies show that few of the relatively smaller punctae were positive for both Atg8-mCherry and GFP (arrowheads), while a majority of the larger punctae were only fluorescent for Atg8-mCherry (white arrows), indicating more acidic punctae (scale bar 10 µm). Genotypes: **(B)** Canton S, **(C)** w^1118^/+;*gl*-TDP-43^CO^/+;+, **(E-H)** w^1118^, GMR-GAL4/ w^1118^;*gl*-TDP-43^CO^/+;UAS-Atg5-GFP/+, **(I-K)** w^1118^, GMR-GAL4/ w^1118^;*gl*-TDP-43^CO^/UAS-Atg8-mCherry-GFP;+.

To further characterize the large vacuoles, we coexpressed CO-TDP-43 and a tagged autophagy protein, Atg5-GFP, which is responsible for the formation of the autophagosomes. We found that these vacuoles were positive for both Atg5 and TDP-43 (Fig. 5E-H). During autophagy, autophagosomes merge with lysosomes to become autolysosomes and are acidified to degrade proteinaceous waste materials (Zhang *et al.* 2013). To determine if the autophagosomes observed were mature and functional autolysosomes, we used the Atg8-mCherry-GFP tandem reporter to assay the relative acidity of the autophagosome/autolysosomes. Atg8-mCherry-GFP is a tandem reporter that detects Atg8, which is localized in autophagic intermediates, and a pH-sensitive GFP that only emits a signal at a neutral pH (Filimonenko *et al.* 2007). This is a useful tool to help understand whether the large vacuoles observed in our CO-TDP-43 flies were autophagic as well as acidic, which is characteristic of autophagic intermediates. We coexpressed CO-TDP-43 with the tandem reporter and found that the larger punctae were positive for Atg8-mCherry, while only a subset of the relatively smaller punctae were stained with GFP, indicating non-acidic compartments (Fig. 5I-K). In contrast, the majority of the punctae that were larger in size were only fluorescent for Atg8-mCherry (Fig. 5I-K), indicating more acidic and mature autolysosomes. These results suggest that misexpression of CO-TDP-43 leads to increased acidic lysosomal vacuoles that are indicative of autophagy upregulation.

## DISCUSSION

To date, very little is known about the exact mechanism of action of TDP-43 mediated toxicity. Here, we report a novel transgenic *Drosophila melanogaster* resource to better understand TDP-43 mediated neurodegeneration. There is great potential for the codon optimized TDP-43 model, as it exhibits robust and sensitive phenotypes ideal for genetic manipulations that allow us to understand its pathogenic mechanisms in an *in vivo* system. Our results suggest that this model has important utility in understanding the TDP-43 mediated pathology in neurodegenerative disorders.

Firstly, while previous studies using fly models helped us understand how the protein leads to toxicity and eventual neurodegeneration, there are several limitations associated with them. The reported models using fly lines overexpressing wild-type TDP-43 did not show strong phenotypes, and conclusions drawn from the fly studies depend heavily upon lines containing pathogenic variants (Lu *et al.* 2009; Li *et al.* 2010; Ritson *et al.* 2010; Voigt *et al.* 2010; Miguel *et al.* 2011; Langellotti *et al.* 2016; Chang and Morton 2017). Similar to published reports, we were previously unable to show any robust phenotypes with the wild-type human transgenic TDP-43 flies (Choksi *et al.* 2014). As a majority of ALS and FTLD cases do not carry known pathogenic mutations, it is critical to understand the mechanism by which the wild-type TDP-43 drives the disease. Compared to previously reported wild-type TDP-43 models, codon-optimized TDP-43 flies exhibit robust eye, wing, and bristle phenotypes, mirroring disease-specific characteristics of TDP-43 (summarized in Table 1). Our findings are in line with previous studies that associated pathogenic mutations in TDP-43 to severely damaged sensory neurons, affecting both the central and peripheral nervous systems in patients (Camdessanche *et al.* 2011). The robust phenotypes observed in our study are indicative of cellular dysfunction and death, and are probable markers for neurodegenerative models. Using genetic and molecular approaches to analyze the mechanisms underlying TDP-43 mediated phenotypes in the eye or the wing may elucidate plausible therapeutic targets of TDP-43.

**Table 1.** Gain-of-function Drosophila models for TDP-43. A comparison of human wild-type TDP-43 transgenic gain-of-function models and codon-optimized TDP-43 models for disease-specific findings.

| **Gain-of-function Drosophila models for TDP-43** |  |  |  |  |  |  |  |  |  |  |  |
| --- | --- | --- | --- | --- | --- | --- | --- | --- | --- | --- | --- |
|  | *CO-TDP-43*<br>*(this paper)* | *Lu et al.*<br>*2009* | *Voigt et al.*<br>*2010* | *Hanson et*<br>*al. 2010* | *Li et al.*<br>*2010* | *Ritson et al.*<br>*2010* | *Estes et al.*<br>*2011* | *Guo et al.*<br>*2011* | *Miguel et*<br>*al. 2011* | *Chang and*<br>*Morton 2017* | *Pons et al.*<br>*2017* |
| Transgenic model | human CO-TDP-43 | human wt TDP-43 | Synthetic wt TDP-43 | human wt TDP-43 | human wt TDP-43 | human wt TDP-43 | human wt TDP-43 | human wt TDP-43 | human wt TDP-43 | human wt TDP-43 | human wt TDP-43 with TBPBR expression |
| Eye phenotype | + | - | - | +<br>(age dependent phenotype) | +<br>(age dependent phenotype) | - | +<br>(at increased temp 29°C) | - | + | - | + |
| Sensory tissue phenotype | + | + | - | - | - | - | - | - |  | - | - |
| Cytoplasmic mislocalization | + | - | - | - | + | - | + | - | + | - | - |
| High-molecular weight/oligomeric species | + | - | - | - | - | - | - | - | - | - | - |
| Cellular aggregation | + | - | - | - | + | - | + | - | - | - | - |
| Autophagic upregulation | + | - | - | - | - | - | - | - | - | - | - |

Secondly, our codon optimized TDP-43 transgenic model affected multiple different cell types in the fly retina, as evident by depigmentation of pigment cells, irregularities in interommatidial bristle cells, disruption in rhabdomeres morphology, disruption of photoreceptor neuron morphology, and necrosis of the cone cells. The effects of neurodegenerative proteins on the Drosophila eye can be diverse. For example, in a polyglutamine-expanded human huntingtin transgenic model, the expanded huntingtin protein was shown to form nuclear inclusions and cause severe degeneration of photoreceptor cells (Jackson *et al.* 1998). Furthermore, the human wild-type tau transgenic model showed abnormal polarity and some rhabdomere loss, mostly affecting the cone cells and ommatidial architecture (Jackson *et al.* 2002; Ambegaokar and Jackson 2011). Based on our observations, TDP-43 pathology is not restricted to photoreceptor neurons, but is most likely widespread among different cell types in the Drosophila retina. In fact, TDP-43 is known to be present and show disease-related pathology across different types of cells both in humans and in animal models (Mackenzie and Rademakers 2008; Wegorzewska *et al.* 2009). Moreover, a Drosophila model of TDP-43 has been shown to exhibit individual responses in motor neurons and glial cells (Estes *et al.* 2011). The codon-optimized TDP-43 flies would therefore serve as an ideal genetic resource to pursue in-depth investigations that determine the morphological effects of TDP-43 in diverse cell types.

Thirdly, in our codon-optimized model, we detected mislocalization of the aggregated form of wild type TDP-43 from the nucleus to the cytoplasm. This phenomenon was validated by the electrophysiological readouts in our study, which showed that toxic aggregates of wild-type TDP-43 reduced functional activity in photoreceptor neurons in the adult eye. The cytoplasmic mislocalization and presence of toxic TDP-43 aggregates have been well characterized in human patient samples of ALS/FTLD (Geser *et al.* 2009; Ritson *et al.* 2010; Miguel *et al.* 2011; Lee *et al.* 2012; Chang and Morton 2017). While *in vitro* studies have shown disease-specific mutant or truncated TDP-43 can form toxic aggregates of oligomeric species, very few studies in wild-type TDP-43 animal models have demonstrated a similar robust production of TDP-43 aggregates (Johnson *et al.* 2009; Couthouis *et al.* 2011; Guo *et al.* 2011; Lee *et al.* 2012; Choksi *et al.* 2014). Our study is unique in that it showed a similar accumulation of toxic aggregates with wild-type human TDP-43 protein. Our codon-optimized lines yield a higher level of TDP-43 protein expression compared to non-codon optimized lines and therefore display more robust toxic phenotypes. This is unsurprising, considering that there have been reports in both sporadic and familial cases of FTLD of increased TDP-43 expression in patient brain tissues (Mishra *et al.* 2007; Gitcho *et al.* 2009). Hence, there is a possibility that TDP-43 has a dosage-dependent effect on its propensity to form toxic aggregates. Several groups have further shown that cytoplasmic mislocalization of TDP-43 causes neuronal toxicity (Shan *et al.* 2009; Barmada *et al.* 2010). Previously, we have been able to show such robust mislocalization of TDP-43 only with disease-specific mutant hTDP-43 Q331K flies (Choksi *et al.* 2014). In keeping with these observations, the disease-specific, dysfunctional phenotypes that we observed with misexpression of wild-type TDP-43 in our codon optimized model offer a great resource to study the cellular processes that could be involved with ALS/FTLD.

Fourthly, in our codon-optimized TDP-43 model, we observed an increase in acidic vacuoles, as evident by lysotracker staining, that are positive for autophagic proteins ATG5 and ATG8 known to be involved in the formation of early and late stage autophagosomes (CITE). As misfolded proteins or toxic protein aggregates are typically cleared by autophagy, a disruption in the cellular process can lead to the accumulation of toxic protein aggregates, which has been linked to many neurodegenerative diseases, including Alzheimer’s disease, Parkinson’s disease, Huntington’s disease, and ALS (Wong and Cuervo 2010; Sasaki 2011). For example, accumulation of autophagosomes was observed in the spinal cord tissues of patients with sporadic ALS (Sasaki 2011). Previously, it has been reported that an inhibition of the ubiquitin proteasome system and autophagy led to increased TDP-43 aggregation and toxicity (Brady *et al.* 2011). In addition, p62, which is a part of the ubiquitin proteasome system, has been identified to directly bind with TDP-43, and its overexpression can reduce TDP-43 aggregation (Tanji *et al.* 2012). Previously, we showed that misexpression of Tau leads to dysfunction of the autophagic process and leads to formation of giant autophagic bodies (Bakhoum *et al.* 2014). Similar to our previous findings, these acidic lysosomal vacuoles observed in codon-optimized TDP-43 model were mature autolysosomes induced to clear the cytoplasmic aggregates of TDP-43. Therefore, upregulation of autophagy in the clearance of TDP-43 proteinopathies could be manipulated as a potential therapeutic target, and our codon-optimized TDP-43 model is an excellent resource for further investigations into this mechanism.

Lastly, TDP-43 is an RNA-binding protein that is involved with RNA metabolism and regulation. As a result, much effort has been devoted to identify the RNA targets of TDP-43 using cell culture models, animal models, and ALS and FTLD patient brain samples. Recently, TDP-43 was shown to bind approximately 30% of the mouse transcriptome, identifying a vast number of possible interactors that can associate with TDP-43 to regulate RNA processing and splicing (Polymenidou *et al.* 2011; Tollervey *et al.* 2011). Many of these putative modifiers bind the UG-rich sequence at introns of TDP-43 (Bhardwaj *et al.* 2013). In this context, our codon-optimized human TDP-43 expressing fly model provides an *in vivo* platform to characterize and validate some of these modifiers to better understand the TDP-43-dependent disease mechanism in ALS/FTLD. In addition, the robust phenotypes observed in the external organs of the eye, wing, and notum of these flies can be scored easily, offering an excellent model for high-throughput screens of modifiers genes that will help elucidate the molecular mechanism of toxicity due to TDP-43. Targeted genetic screens that identify effectors of TDP-43 will allow us to further identify and pursue novel mechanisms for disease pathology.

## ACKNOWLEDGEMENTS

We would like to thank Dr. Fen-Biao Gao for providing us with the human wild-type TDP-43 transgenic fly stock and Dr. Hugo J. Bellen for the *eq*-GAL4 stock. We would like to thank Dr. P. Robin Hiesinger for the GMR-RFP stock and for allowing us to use his laboratory and equipment to perform electroretinogram experiments, as well as Daniel Epstein for his help with the experiments. We would also like to thank Dr. Santhosh Girirajan, Dr. Suren Ambegaokar, Matthew Jensen, and Vijay Kumar for their helpful discussions and comments.

